# Dissection of the contribution of bushy cell delays to sound localization in juvenile and adult Mongolian gerbils

**DOI:** 10.64898/2026.10.30.762924

**Authors:** M. C. Sierksma, J. G. G. Borst

## Abstract

Sound sources along the azimuth generate interaural time differences in arrival (ITDs) at the ears. According to Jeffress’ hypothesis, ITDs are detected by coincidence-sensitive neurons of the medial superior olive (MSO) for which a differential axonal delay from each ear to the MSO compensates for the ITDs. The origin and development of these delays remain poorly understood. To this end, we made recordings from bushy cells (BCs), which project to both MSOs, in adult and in 2-4 week old Mongolian gerbils. First spike latency revealed a frequency-dependent delay likely originating in the cochlea. Latencies were markedly larger for pure tone stimulation than for complex sounds. Combined with our published recordings from the MSO, we observed a developmental decrease in most delays, including the contralateral BC-to-MSO axonal delay. Unexpectedly, the ipsilateral BC-to-MSO axonal delay did not decrease. Anatomical brainstem reconstructions revealed that the inter-MSO distance was stable whereas the cochlear nucleus-to-MSO distance increased during development. Together, this indicated an increase in axonal propagation velocity, particularly for the contralateral projection. The variability in the averaged firing onset between pairs of BCs was found to be as large as the variance in bITDs, suggesting that BCs with matching firing onset are selected to converge onto the same MSO neuron. Together, our results indicate that the developmental speedup of ear-to-MSO delays contributes greatly to bringing the bITDs within their ecological range, but that the difference in path length is not entirely overcome, keeping the ITD tuning biased to contralateral sound sources.

**Significance statement:** Humans determine the spatial origin of sounds by comparing their arrival time at either ear. Medial superior olive (MSO) neurons read out these sub-millisecond differences, despite responding ∼7 ms after sound onset. How such remarkable precision is achieved remains unanswered. We dissected the sources of delay and timing variability along the pathway from ear to MSO. Even though the contralateral axonal delay travel speeds up more than the ipsilateral during development, MSO responses remained biased toward contralateral sounds. While classical models emphasize axonal delays as the source of internal timing differences, large variability arises before signals reach the MSO-projecting axons, which necessitates a developmental selection mechanism to overcome this variability and improve the sharpness of ITD tuning.

## Introduction

Interaural differences in arrival time at the ears (ITD) are perceived as a sound coming from a specific direction. ITDs are detected by the principal neurons of the medial superior olive (MSO). They receive excitatory innervation from a limited number of spherical bushy cells located in both the ipsi- and the contralateral anteroventral cochlear nuclei (AVCN; (Osen, 1969; Smith et al., 1993; Beckius et al., 1999; Couchman et al., 2010; Karino et al., 2011; Franzen et al., 2020; Callan et al., 2021). These bushy cells (BCs) fire with high temporal precision (Joris et al., 1994b; Kuenzel et al., 2011; Keine and Rübsamen, 2015), allowing their EPSPs to preserve fine temporal features of sound. Coincidence of ipsi- and contralaterally evoked EPSPs determines MSO firing and underlies ITD tuning (van der Heijden et al., 2013). Individual MSO neurons differ in the ITD evoking the maximal firing rate, *i.e.* the best ITD (bITD). At the bITD there is maximal coincidence of EPSPs, as the difference in ear-MSO latencies between ipsi- and contralateral pathways compensates for the ITD. This difference in pathway latency is called the internal delay (Jeffress, 1948; Goldberg and Brown, 1969; Yin and Chan, 1990; Spitzer and Semple, 1995; van der Heijden et al., 2013; Plauška et al., 2016; Mackenbach and Borst, 2023; Sierksma and Borst, 2024).

The Jeffress’ hypothesis postulates that the internal delay arises from the difference in axonal lengths between BCs (Jeffress, 1948); a systematic arrangement of axonal length differences to create a spatial map of ITD tuning was postulated, but found no anatomical support in mammals (Smith et al., 1993; Beckius et al., 1999; Karino et al., 2011; Franken et al., 2015). Alternatives for the nature of the internal delay have been put forward, including stereausis (Shamma et al., 1989), well-timed inhibition (Brand et al., 2002; Pecka et al., 2008; Grothe et al., 2010), differences in the dendritic trees and axonal location of MSO neurons (Jercog et al., 2010; Winters et al., 2017; van der Heijden, 2018; Callan et al., 2021; Mackenbach and Borst, 2023; Sierksma and Borst, 2024). Some hypotheses have received limited support in mammals (Roberts et al., 2013; van der Heijden et al., 2013; Franken et al., 2015; Plauška et al., 2016; Plauška et al., 2017; Mackenbach and Borst, 2023; Sierksma and Borst, 2024), but, alternatively, the nature of the internal delay may be cumulative and complex, deriving from many delays encountered along the pathway.

One approach to further delineate the mechanisms that determine ITD tuning is to study the developmental changes in the brainstem pathways leading up to the MSO. In our previous work we studied the developmental changes in ITD tuning of MSO neurons. We found that shortly after hearing onset the MSO neurons were tuned to contralateral sounds and that their bITD was restricted to a relatively small range that was well outside of the ecological range. In the next weeks the bITDs gradually shifted towards the ecological range. We showed that internal delay may be the outcome of a developmental mechanism selecting synaptic inputs of similar latencies (Sierksma and Borst, 2024). At the same time, we observed that the ear-MSO latency decreased after hearing onset. Most, if not all, of the auditory brainstem delays continue to mature after hearing onset (Seidl and Rubel, 2016; Stange-Marten et al., 2017; Anbuhl et al., 2025). Together with the maturation of the intrinsic properties of the bushy cell (Müller et al., 2019) and the endbulb synapse (Bellingham et al., 1998) this creates a moving target for selecting inputs based on latencies.

The debate on the emergence of ITD tuning could be advanced if the contributions of individual delays along the neural pathway toward the MSO neuron were elucidated. Currently, little is known about these individual delays underlying the ear-MSO latency nor how they develop after hearing onset. To determine how individual delays along the pathway contribute to the internal delay, we recorded from bushy cells from both postnatal day (P)15-28 and adult gerbils and combined these measurements with our published MSO latencies (Sierksma and Borst, 2024). This allowed us to test whether the developmental change in ear-MSO delay can be assigned to the delay arising along the bushy axon, by synaptic integration or to early processing stages. Furthermore, we compared the variability in bushy latencies with the ecological range, which will be discussed in the context of the Jeffress’ model.

## Methods

### Animals

All experiments were conducted in accordance with the European Communities Council Directive (86/609/EEC), and were approved by the national and local ethical committees.

Breeding pairs of Mongolian gerbils (*Meriones unguiculatus*, Janvier Labs) were kept in the animal facilities of the Erasmus MC. The gerbils were given ad libitum water and food. When the female appeared pregnant, the cage was checked daily. Postnatal day (P)0 was defined as the day the litter was first observed.

### Surgery

The surgical approach for young gerbils was adapted from a published procedure for adult gerbils (Frisina et al., 1982). Juvenile Mongolian gerbils were anesthetized using an intraperitoneal (i.p.) injection with a mix of ketamine/xylazine (80/15 mg/kg, respectively). Additional i.p. injections with a ketamine/xylazine mix (adult: 16/3 mg/kg, juvenile: 10/0.2 mg/kg) were given when the hind paw reflex started to return. Rectal temperature was maintained at 36.5 to 37.5°C using a homeothermic blanket (Stoelting Co.). The bony external meatus was exposed to obtain an unobstructed opening to the eardrum. A metal plate was glued to an exposed rostrodorsal part of the skull to fix the head. The animal continued to breathe independently, and was euthanized if breathing became disturbed. Soft tissue covering the dorsocaudal bulla was removed and a small hole was made using a sharp needle and forceps. Then, a small hole was cut using a sharp needle from the temporal bone covering the paraflocculus. A reference electrode was inserted in the vermis of the cerebellum.

### Electrophysiology

The rostral AVCN was targeted to record from low-frequency bushy cells, presumably the spherical bushy cells (Frisina et al., 1982; Kuenzel et al., 2011). As other cochlear neurons were also encountered in the rostral AVCN, we rejected post-hoc any recording in which there was no presence of a prepotential, which is a hallmark of the giant endbulb of Held terminal contacting the bushy cell (Pfeiffer, 1966; Kopp-Scheinpflug et al., 2002; Englitz et al., 2009; Kuenzel et al., 2011; Keine and Rübsamen, 2015; Müller et al., 2019). Loose-patch (“juxtacellular”) or unit recordings were made from the cochlear nucleus neurons using thick-walled borosilicate glass pipettes with filament. The pipette had an open tip resistance of 5 to 8 MΩ when filled with NaCl 135, KCl 5.4, CaCl_2_ 1.8, MgCl_2_ 1, HEPES 5 mM adjusted to pH 7.2 with 1 M NaOH. On the day of recording, the pipette solution was sometimes supplemented with 1-2% biocytin. The recording electrode was oriented vertically (young) or slightly angled toward rostral (adult), and lowered through the paraflocculus. Click-evoked field potentials and spiking activity were used to assess the entry into the cochlear nucleus. The recording pipette was lowered into the brain under high positive intra-pipette pressure (>100 mbar) until entering the cochlear nucleus (30 mbar). The pipette was slowly advanced until the resistance would slightly increase and then pressure was reduced to 0-5 mbar. The pipette was advanced until action potentials appeared. Recordings were performed in current-clamp mode with bridge balance and pipette capacitance compensation. Data were acquired using a MultiClamp 700B amplifier (Molecular Devices), low-pass filtered at 10 kHz with a four-pole Bessel filter, and sampled at 111.6 kHz using custom software (courtesy of Dr. Van der Heijden) written in Matlab 7.6.0 (Mathworks).

If the animal was not euthanized at the end of the session, it was deeply anesthetized and transcardially perfused with saline and 4% paraformaldehyde, and the brain was removed for further processing.

### Auditory stimuli

#### Setup

Auditory stimuli have been described in Sierksma and Borst (2024). Briefly, auditory stimuli were generated using custom MATLAB software and realized through a 24-bit DA processor (RX6, Tucker Davis Technologies (TDT), Alachua), a programmable attenuator (PA5, TDT), and an amplifier (SA1, TDT). Sounds were presented to the animal in a close-field configuration with Shure speakers (22 Hz to 17.5 kHz) that were each connected to a metal bar positioned in front of the external meatus. Click intensity was calibrated using a 1/4” condenser microphone (model 7016, ACO Co). Four oscillations were observed in the first millisecond of the click. The maximum of the peak-to-peak amplitudes of these four oscillations was measured using an oscilloscope and converted to sound pressure level based on the sensitivity of the microphone.

#### Monaural clicks

We used monaural clicks, a 0.1 ms condensation pulse, with an interval of 30 ms between contralateral and ipsilateral, repeated every 200 ms with varying amplitudes (click-intensity protocol) in the range of 0-60 dB SPL with steps of 3-6 dB. The number of repetitions ranged from 15 to 30.

#### Frequency-response area

We measured the frequency response area of cells using pure tones ranging from 0.5 to 7.5 kHz and intensities from 0 to 80 dB SPL. Starting sine phase was 0 for each tone. Tone duration was 50 ms including a 3 ms cos^2^ on/off ramp. A tone was presented every 500 ms. The Frequency-SPL set was repeated three times.

#### Zwuis-intensity

The wideband multitone Zwuis (van der Heijden and Joris, 2003) ranged between 100 and 5,000 Hz with 45 tone frequencies and random start phases for each frequency. These start phases were kept the same per animal. The amplitude of each frequency component was equal, and the stimulus intensity ranged from 0 to 50 dB SPL. Burst duration was 300 ms including a 10 ms cos^2^ ramp at its on- and offset. It was presented every 600 ms. Number of repetitions ranged from 4 to 10.

For all stimuli, presentations were randomized per repetition.

### Histology and widefield imaging

The perfused brains were extracted and fixed for 2 hr at room temperature in 4% paraformaldehyde or overnight at 4 °C. The brains were cryoprotected overnight in 10% sucrose in 0.1 M PB, embedded in 14% gelatin/10% sucrose, fixed in 10% formaldehyde/30% sucrose for 3h at room temperature, and stored in 30% sucrose in 0.1M PB at 4 °C for at least two days. Coronal sections (50 μm) were cut on a freezing microtome and rinsed three times in 0.1 M PB. Next, the sections were cooked at 70-80 °C for 3h in 10 mM sodium citrate (pH 6), rinsed three times with 0.9% (w/v) NaCl in 0.05 M PB (phosphate-buffered saline; PBS). The sections were then pre-absorbed for 1h with 10% (v/v) normal horse serum and 10% (v/v) Triton X-100 in PBS followed by 36–48 h incubation at 4 °C under gentle agitation. The incubation solution contained 2% (v/v) normal horse serum, 0.4% (v/v) Triton X-100 and the primary antibodies. The slices were washed three times in PBS, and then incubated under gentle agitation either for 2h at RT or overnight at 4 °C in the secondary antibody solution containing 2% (v/v) normal horse serum, 0.4% (v/v) Triton X-100, and the secondary antibodies. The sections were washed once with PBS, and incubated for 10 min RT in 0.3 µM 4′,6-diamidino-2-phenylindole (DAPI) (D3571, Thermo Fisher Scientific) in 0.1 m PB. Sections were then washed three times with 0.1 M PB, mounted on glass coverslips with gelatin-chrome alum, air-dried, and closed with Mowiol mounting solution containing 10% (w/v) Mowiol 4–88, 25% (v/v) glycerol in 0.1 m Tris-buffer (pH 8.5). Sections were kept at 4 °C in the dark until further use.

The primary antibodies were guinea-pig polyclonal anti-vesicular glutamate transporter 1 (Millipore, Burlington, MA, USA, AB5905, RRID: AB_2301751; 1:3000) and 2 (Millipore, AB2251, RRID: AB_1587626; 1:3000), or sheep polyclonal anti-glycine transporter 2 (Millipore, AB1771, RRID: AB_909451:2000), or all three. Secondary antibodies were Alexa Fluor 555 against sheep and Alexa Fluor 488 or 645 against guinea pig (1:200, Jackson ImmunoResearch, Ely, UK).

Sections were imaged using an Axio Imager 2 (Zeiss, Germany), equipped with an Axiocam 705 camera (Zeiss, Germany), 5×/0.16 NA objective and optimized acquisition settings.

### Electrophysiological analysis

All analyses were done using custom-written codes in Matlab R2024b (MathWorks).

#### Event detection

Extracellular action potentials were detected offline based on a manually set threshold for the maximum repolarization rate of individual events. Only cells for which histograms of the negative side of the first derivative showed bimodality were accepted for further analysis (van der Heijden et al., 2013).

#### Pure tone responses

For each Frequency-SPL combination, the firing rate during sound presentation was calculated and the average firing rate was plotted as a heat map. The characteristic frequency (CF) was visually identified. Tone-evoked first spike latencies were corrected for background firing as follows: we created a cumulative distribution of the number of spikes following the onset of sound, this distribution was compared to the expected number of spikes from the spontaneous activity of the unit + 1 spike to identify the time at which the first sound-evoked spike occurred. This timepoint was taken as the first spike latency (FSL), and it was calculated independently for each tone-intensity combination and repetition. We only used the four highest intensities to calculate the average FSL.

Group delay was estimated from the responses to tone stimulation. Average phase was calculated per tone presentation for which the vector strength was >0.3. Vector strength was calculated as described by Goldberg and Brown (1969). The wrapped phase-frequency was fitted per intensity using φ(f) = φ(0) −2πfτ, with τ as the group delay (FitzGerald et al., 2001). A grid search from 1 to 20 ms for τ was performed to find the minimal summed squared circular error. For most adult bushy cells we observed a periodicity of 2 ms in the best group delay, meaning that there was a family of best group delays (τ + n*2 ms) for each cell. Across intensity we took the median to obtain the group delay for the bushy cell. The group delays were usually ∼13 ms. Group delays less than 13 ms were corrected by multiples of 2 ms to have a similar group delay.

#### Click latency

eAPs from +2.5 ms to +10.5 ms following the click were selected. The median latency of this selection was calculated. The selected eAPs within less than 1 ms of the median latency were selected, and the median latency of this subset was taken as the FSL to the click. Next, the median latency was calculated for each click amplitude. To quantify the shift in latency at increased sound intensities, we calculated for each neuron the difference between the FSL and the median latency for the click with the smallest click amplitude for which we recorded >3 eAPs. To measure the cochlear delay we fitted the median latency with a power relation against CF (Versteegh et al., 2011): FSL = 3.53 ms + 0.96 CF^-0.812^ (CF in kHz). When we write that we corrected for cochlear delay in the paper, we subtracted 0.96 CF^-0.812^.

#### Zwuis responses

Spikes during the presentation of Zwuis stimuli were used for the following calculations. For each intensity we calculated the average firing rate. For each intensity and frequency the vector strength and average phase of the spike times were calculated. For eight bushy cells, phase-locking was highest for a frequency below 300 Hz and higher than to the higher CF. Consequently, we excluded frequencies below 300 Hz from further analysis. To identify the stable-delay frequency (sdF), the standard deviation of the phase across intensities was calculated per frequency. The frequency with the smallest standard deviation was taken as sdF. The phase of this frequency was transformed to a delay by multiplying the phase with the frequency period. To allow a better comparison between cells of how the delay depends on intensity, we subtracted the cell’s average phase/delay at this stable-delay frequency across intensities.

#### Complex waveform analysis

A snippet of the recording was extracted ranging from -5 to +4 ms centered at spike peak for each spike during the Zwuis stimuli and the inter-stimulus periods. The EPSP onset was calculated as before (Sierksma and Borst, 2024). Briefly, the EPSP onset was defined by a threshold crossing from the start of the snippet. This threshold was equal to the median value of the snippet plus an offset which was taken as the lowest value of either 10% of the spike amplitude or two times the standard deviation of the snippet 3 to 0.5 ms prior to the spike peak. The EPSP max rate of rise (ror) was defined as the peak in the first derivative of the snippet at timepoints later than the EPSP onset and less than 0.15 ms before the spike peak. The EPSP-AP delay was calculated as the time difference between EPSP onset and spike peak. To compare EPSP max ror and EPSP-AP delay between cells, we normalized the EPSP max ror to the average max ror for ISI >30 ms and we subtracted the EPSP-AP delay with the average EPSP-AP delay for ISI >30 ms. The prespike-spike delay was visually defined by overlaying the snippets and manually indicating the prespike peak. We did not attempt to identify prespikes for single snippets due to the small amplitude of the prespike for some recordings.

#### MSO data

MSO latencies, CFs and EPSP-AP delays were taken from our previous publications (Plauška et al., 2017; Sierksma and Borst, 2024). These data were collected on the same setup using similar stimulation protocols.

### Anatomical analysis

Widefield images were first checked for their proper order and orientation. Their orientations were aligned by a manually drawn midline, resulting in rotating and shifting the image. Then, we manually outlined the AVCNs and MSO for each image. AVCN was delimited caudally by the auditory nerve entry. The MSO was identified by its GlyT2-positive/VGLuT-negative somatic layer sandwiched between GlyT2-negative/VGLuT-positive dendritic areas. The somatic layer of the MSO was also indicated by drawing a ventrodorsally oriented line segment that followed the somatic layer. The outlines were filled, followed by sequentially registering the outlines with the next image, keeping the middle image at a fixed position. Image registration included only a vertical shift as the midline was already defined per image. Volume rendering was done using 25×25×25 μm-voxels. Each outline was resampled to contain 500 points and interpolated in rostrocaudal direction by connecting these points to the corresponding points of the outline in the next image.

To measure the dimensions of the nuclei, we created a bounding box with three lengths. These bounding boxes were not perfectly aligned to the anatomical directions (rostrocaudal (RC), ventrodorsal (VD) and mediolateral (ML)). The length of the side best fitting to the anatomical direction was used to indicate the size of the nucleus in this anatomical direction. Volume of the nucleus was taken as the number of positive voxels.

To measure inter-MSO distances, we quantified the average nearest-neighbor Euclidean distance from each positive voxel of the MSO somatic layer to the positive voxels of the contralateral MSO somatic layer. To measure cochlear nucleus (CN)-MSO distance, we quantified pair-wise Euclidean distances from positive CN voxels to positive ipsilateral MSO somatic voxels. The average was taken as the distance between CN to iMSO.

### Experimental design and statistical analysis

Our main design was to describe developmental trends by making the same measurements at different ages. Mean and sample standard deviations are reported as mean ± SD. Pearson’s correlation coefficient (*r*) was calculated for linearly related variables, and Spearman’s correlation coefficient (*rho*) when variables were related by a power function, sigmoid function, or when the variables were correlated against age including the adult data. Data were fitted using Matlab functions *fitlm* (linear regression) or *fitnlm* (non-linear regression). P-values indicate the *F*-test of the model. To test for a difference between fit coefficients for the same model, we used a z-test. To test for a difference in propagation velocity, we used a paired *T*-test. Empirical cumulative distributions were tested using a two-sample Kolmogorov-Smirnov test.

## Results

### Identification of bushy cells in young and adult gerbils

In our previous study we described the neural latencies for MSO activity (Sierksma and Borst, 2024). These ear-MSO latencies are determined by the bushy cell latency (ear-CN) plus a delay between bushy cells (BCs) and MSO cells (CN-iMSO, CN-cMSO, Figure 1A). To dissect these latencies (Figure 1A), we made recordings from the CN for a total of 73 and 37 cells from 18 young (postnatal day P15-P28) and 10 adult, anesthetized Mongolian gerbils, respectively. Biocytin labeling confirmed that the recordings were obtained from the anteroventral cochlear nucleus (CN, Figure 1B). For none of the cells for which we also performed contralateral stimulation, cell’s firing correlated with contralateral sound presentation or was modulated by ITDs. The recorded cells were thus monaurally tuned.

**Figure 1.**
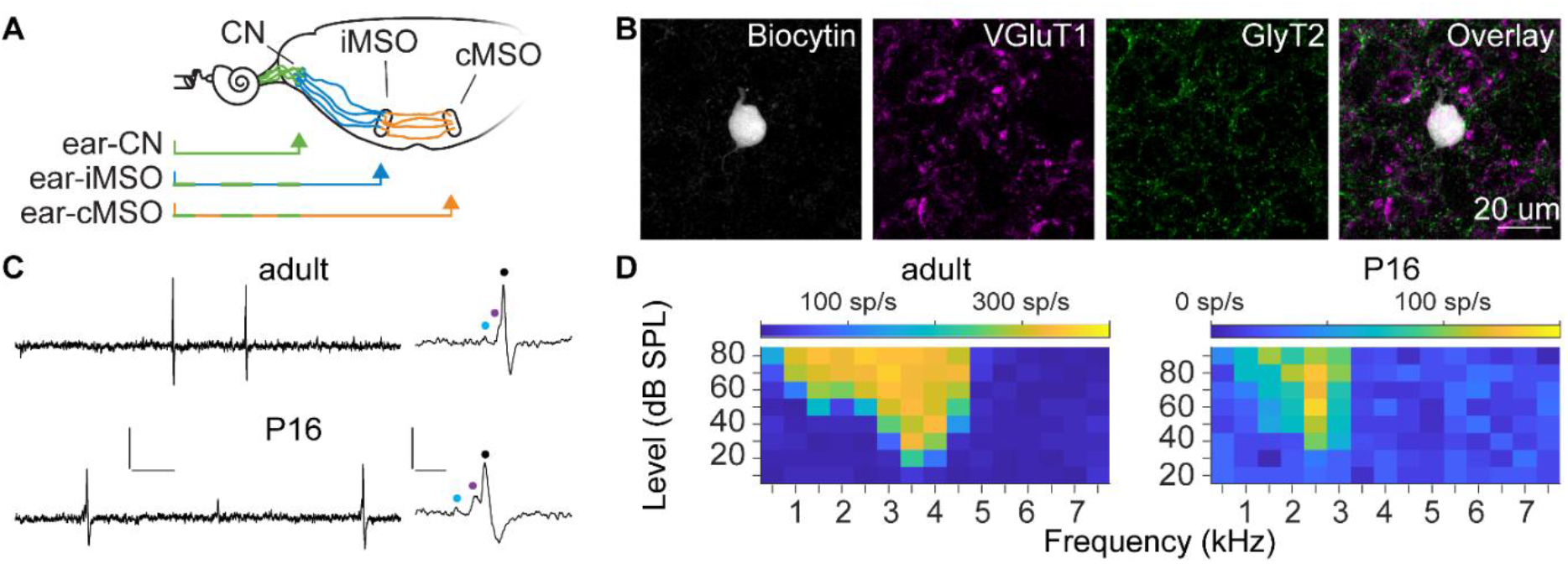
Identification of bushy cells in vivo. (A) Schematic drawing of the auditory brainstem. Neural delays to the MSO include a delay from ear to cochlear nucleus (CN, green arrow) and an ipsi- and contralateral delay from CN to MSO (blue and orange, respectively). (B) From left to right, Biocytin labelling of a bushy cell (left), excitatory (VGluT1) and inhibitory inputs (GlyT2), and an overlay. (C) Left, representative recordings of spontaneous bushy activity from an adult (top) and a P1c gerbil (bottom). Right, a complex waveform shown on an expanded time scale, including a prespike (blue marker), eEPSP (magenta marker) and eAP (black marker). (D) FRAs of an adult and a P1c bushy cell. Scale bar in C: 1 mV, 10 ms or 1 mV, 1 ms.

The electrical activity of BCs displayed a complex waveform containing a prespike, an eEPSP and an eAP (Figure 1C)(Pfeiffer, 1966; Kopp-Scheinpflug et al., 2002; Englitz et al., 2009; Müller et al., 2019). Prespikes were observed for 52 of 73 cells and 25 of 37 cells in young and adult gerbils, respectively. From hereon we refer to cells with a prespike as BCs. These cells often showed failures during spontaneous activity (Figure 1C)(Pfeiffer, 1966; Kopp-Scheinpflug et al., 2002; Englitz et al., 2009; Kuenzel et al., 2011; Keine and Rübsamen, 2015). For 40 young cells and 17 adult cells we obtained their frequency-intensity response areas (FRA), characterized by a clear widening below the characteristic frequency (CF) but not above (Figure 1D). Inhibition was often recruited at frequencies above CF. The CF of bushy cells was 2.5 ± 1.3 kHz (min-max, 0.5-5.5) and 2.9 ± 1.9 kHz (0.5-7.5) for young and adult gerbils, confirming that we targeted the rostral anteroventral CN, where the spherical bushy cells are located (Kuenzel et al., 2011; Mylius et al., 2013; Keine and Rübsamen, 2015).

Spherical bushy and MSO neurons are typically tuned to low frequencies (Kopp-Scheinpflug et al., 2002; Pecka et al., 2008; Englitz et al., 2009; Kuenzel et al., 2011; van der Heijden et al., 2013; Franken et al., 2015; Plauška et al., 2017; Sierksma and Borst, 2024). At these frequencies the auditory system can encode the phase of the sound. The accuracy of phase encoding can be quantified as vector strength (Goldberg and Brown, 1969). To measure the frequency with the highest vector strength (i.e. to which the BC phase-locked best), we presented the multitone Zwuis stimulus (van der Heijden and Joris, 2003; Plauška et al., 2016). BCs typically responded to a few frequencies within the Zwuis, allowing us to identify the frequency with the highest vector strength. This frequency was 1.77 ± 0.78 kHz (n = 41 cells, min-max, 0.50-3.93 kHz) and 1.32 ± 0.80 kHz (n = 22 cells, min-max, 0.39-3.02 kHz) for young and adult gerbils, respectively. For 16 adult BCs and 34 young BCs we measured both CF and the frequency to which they phase-locked best. Ten BCs had a CF >3 kHz. At such high frequencies phase encoding degrades (Versteegh et al., 2011; Wei et al., 2017) due to the hair cell’s membrane time constant (Palmer and Russell, 1986). Their CF was not correlated with their best phase-locking frequency (*r* = −0.50). In contrast, CF was strongly correlated with the best phase-locking frequency for the other 40 cells (*r* = 0.86, Freq_best VS_ = -0.11 kHz + 0.91 CF). These findings confirm that we recorded from BCs that encode phases which underlie the ITD responses of the MSO neurons.

### Timing of low-CF bushy firing is level-invariant and independent of tones

To study the timing of bushy firing, we first analyzed the first spike latencies (FSLs) for pure tones and clicks in adult gerbils. These latencies showed an inverse relationship with the frequency tuning of the BC due to the cochlear delay (Figure 2A-B). The click-induced latency followed this relationship (*rho* = –0.8, *p* = 3 10^-8^, FSL = 3.53 ms + 0.96 CF^-0.812^) and was in line with the cochlear delay measured at the auditory nerve (Versteegh et al., 2011).

**Figure 2.**
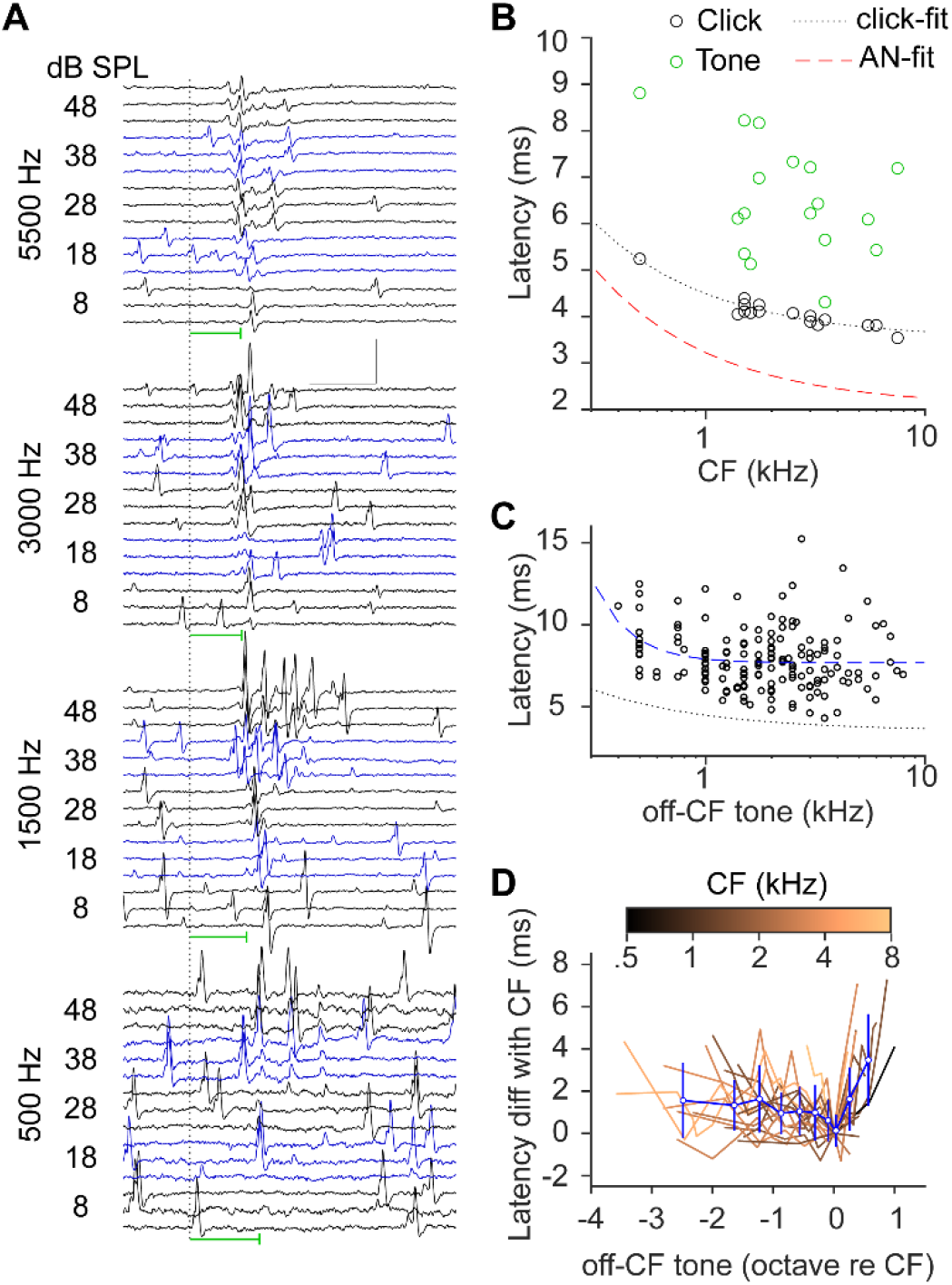
Cochlear delay at bushy cells. (A) Click responses of bushy cells. Responses of four example cells are shown ordered by descending CF (top-bottom). For each cell, responses were elicited by clicks of increasing intensity (bottom-top); per intensity three examples are shown (same-colored traces). Vertical dotted line indicates click onset. Horizontal green line indicates median latency of the cell. Scale bar: 4 mV, 5 ms, (B) Median latency against CF. Black markers are latencies obtained with clicks, green markers are latencies obtained with pure tones. Click latencies were fitted with a power function: FSL = 3.53 ms + 0.Sc CF^-0.812^. Red dashed line: the fit by Versteegh et al. for the auditory nerve shown for comparison: FSL = 2.07 ms + 1.15 CF^-0.808^. (C) Latencies obtained with non-CF pure tones against the tone frequency. A power fit was made: FSL = 7.c8 ms + 0.22 CF^-2.c3^ (blue dashed line). For comparison, the click latency fit is shown (black dotted line). (D) Latency difference with CF for off-CF pure tone stimulation against the tone frequency relative to CF. Each line corresponds to a bushy cell and its CF is color-coded. The expected latency difference based on the click latency fit is shown as a dashed line.

The latencies observed with pure tone stimulation were larger and more variable between BCs (*rho* = –0.41, *p* = 0.1, Figure 2B)(Typlt et al., 2012). This variation obscures the cochlear delay. Off-Cf tone stimulation also creates a delay, observed at the level of the auditory nerve (Versteegh et al., 2011), and below-CF stimulation triggers increased phase-locking, higher entrainment and sometimes peak splitting in bushy cells (Joris et al., 1994a; Recio-Spinoso, 2012; Wei et al., 2017). Importantly, the timing appears less well defined (Wei et al., 2017), which has consequences for ITD computation. We observed that FSLs were larger when stimulated with off-CF tones than expected from the click latencies (Figure 2C-D). To estimate the cochlear delay from off-CF stimulation, we performed a power fit on the data (*rho* = –0.18, *p* = 0.02, Figure 2C) of which only the offset coefficient was significantly different (z = 15.0, p < 0.001). The cochlear delay estimate was thus similar. Still, off-CF stimulation included a delay relative to clicks.

A striking observation was the variation in delay between BCs with a similar CF that were stimulated with off-CF tones. It was in the order of ms, which is large compared to the ecological range of Mongolian gerbils (±0.13 ms (Maki and Furukawa, 2005)). The poor timing of BCs to tones suggests that BCs are not well synchronized during tonal stimulation. As MSO neurons are sensitive coincidence detectors (Mathews et al., 2010; van der Heijden et al., 2013; Mackenbach and Borst, 2023), the non-synchronized BC activity during pure tones may considerably alter the ITD tuning of MSO neurons considering that the off-CF delay would not necessarily be equal for the ipsi- and the contralateral side. We conclude that, contrary to click stimulation, the first spike of bushy cells activated by pure tones includes an unpredictably large, extra delay.

To obtain a delay from the ongoing activity to a tone, we calculated the phase at which an AP occurred during sound stimulation. These phases were then used to fit the group delay (FitzGerald et al., 2001). These group delays were 13.8 ± 0.6 ms (min-max: 12.9-15.0 ms) and were negatively correlated with CF (rho = −0.72). The group delays were well fitted by a linear function, giving another estimate for the cochlear delay (group delay = 14.4 ms − 1.7 ms log_10_(CF/1 kHz)). Overall, group delays were larger than the FSL, but nonetheless shaped by CF. Moreover, the FSLs and the group delays were correlated (group delay = 8.9 ms + 1.2 FSL, r = 0.72). Using these three fits for the cochlear delays, we estimate a latency difference between 4 kHz and 0.5 kHz of 1.38, 1.38 and 1.67 ms from the FSL, off-CF FSL and group delays. The cochlear delay thus shapes the timing of bushy activity, despite the disparate offsets for the different stimulation paradigms, and it contributes up to a ∼1.5 ms difference in latency between adult bushy cells.

Another essential feature of bushy cell firing is that its timing needs to be invariant over time and across intensities. We first assessed how the timing of bushy cell firing changed with sound intensities. Indeed, bushy cell’s FSL depended little on sound intensity, as it decreased only 0.20 ± 0.34 ms with increasing click intensities (threshold to max. test intensity: 36 ± 14 dB), independent of the cell’s CF (*r* = –0.15). To assess the intensity effects for ongoing sounds, we analyzed Zwuis at different intensities for 18 adult bushy cells (Figure 3A). Bushy firing was often phase-locked to some of the Zwuis frequencies with a remarkably stable average phase across 5-50 dB SPL (Figure 3A-B). Bushy cells were typically inhibited at higher levels (9 out of 18), likely due to inhibition recruited at higher intensities (Kuenzel et al., 2011; Nerlich et al., 2014; Keine and Rübsamen, 2015), but they could also show increasing or plateau-like intensity tuning (Figure 3C). Temporal dispersion during stimulation was in the order of 0.1 ms (Joris et al., 1994b; Joris et al., 1994a; Paolini et al., 2001; Recio-Spinoso, 2012). This is often analyzed as vector strength (Dehmel et al., 2010; Kuenzel et al., 2011; Keine and Rübsamen, 2015), but vector strength does not reflect whether the phase/timing is stable across intensities. Instead, we obtained the frequency with the least variation in phase across sound levels. The frequency-phase combination is equal to a delay from which we identified the stable-delay frequency (sdF). SdF was correlated to the frequency with the highest vector strength (r = 0.83). The difference in delay across levels ranged from 0.04 to 0.20 ms (Figure 3D), excluding two outliers (0.60 and 0.67 ms), demonstrating the level-insensitivity of the timing of bushy cells. (The sound envelope is not changing during the Zwuis, which may explain the difference with tone-in-noise experiments (Gai and Carney, 2006)). For 13 of 18 cells we obtained both CF and sdF (Figure 3E) and they were correlated for CF <= 3.0 kHz (*r* = 0.92, sdF = 0.16 kHz + 0.70 CF). The sdF of the three cells with CF >3 kHz was below CF and these cells included the two outliers for sdF. Bushy cell firing for CF <3.5 kHz thus remains well-timed to its CF in ongoing complex sounds across a range of sound intensities.

**Figure 3.**
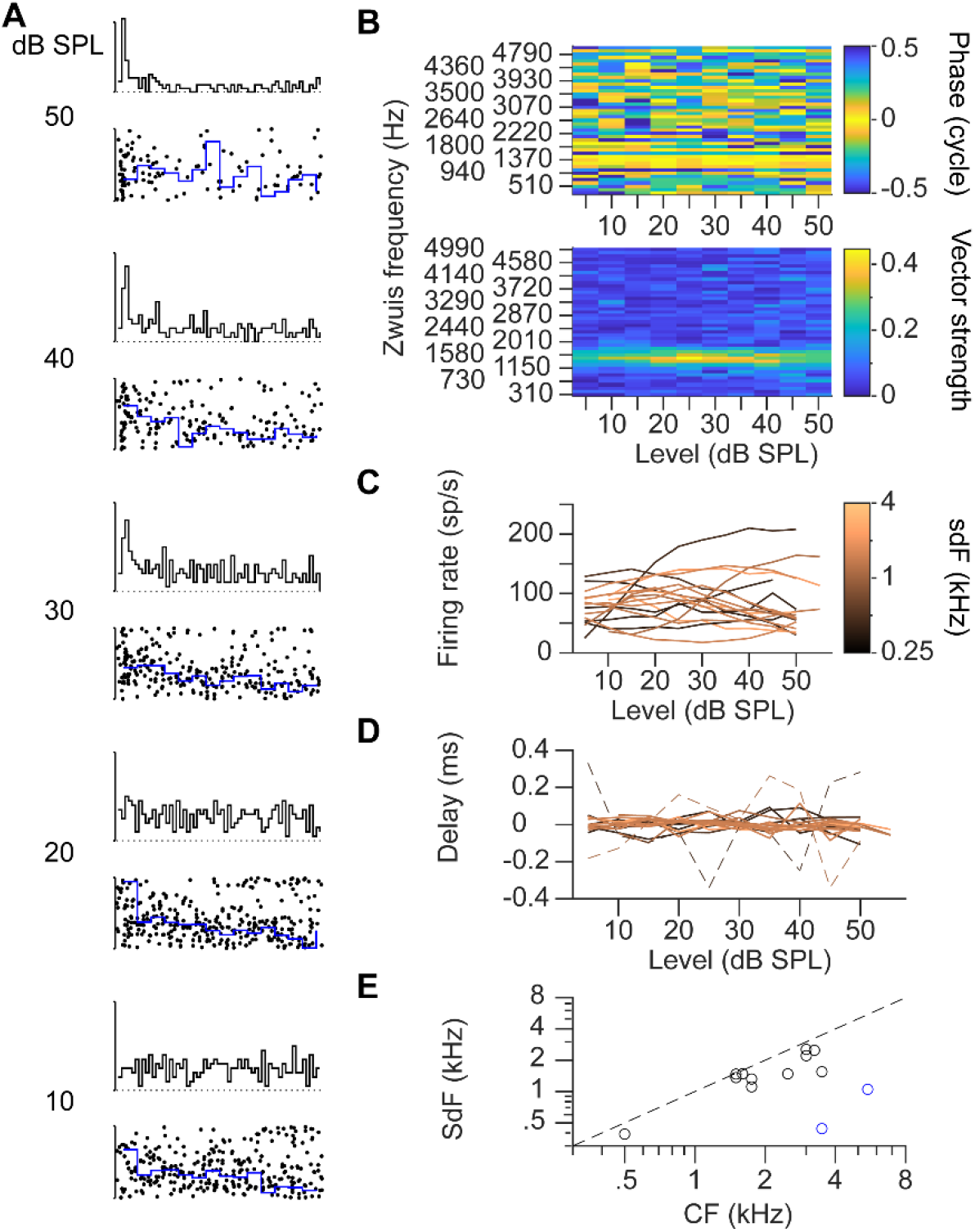
Level-independent timing of bushy cells. (A) Example of firing rate and phase during Zwuis stimulation at different levels in a bushy neuron. Phase-stable frequency is 1.4S kHz. Duration of the Zwuis is 300 ms. Firing rate full scale is 400 spikes/s, 5 ms bins. Dotted line indicates 0 spikes/s. Phase of each AP relative to 1.4S kHz (black dot) and moving average of phase (blue line, 20 ms bins) are shown below each firing rate plot. Phase scale is 1 cycle. (B) Heat map of average phase (top) and vector strength (bottom) per Zwuis frequency of the same cell in A. (C) Firing rate against Zwuis level. Each line corresponds to a bushy cell. Color corresponds to the stable-delay frequency of the bushy cell (sdF). (D) Delay at sdF against Zwuis level. Each line corresponds to a bushy cell color-coded by sdF as in C. In each cell the average delay at sdF has been subtracted. Dashed lines are two outliers. (E) SdF against tonal CF. Each marker corresponds to a bushy cell. Blue markers are the outliers in D.

### Timing of first spike and ongoing activity of developing bushy cells

The ear-MSO latency decreases in the two week period following hearing onset (Sierksma and Borst, 2024). We investigated whether this decrease in latency originates before or after bushy cells, or both. Recordings from young Mongolian gerbils showed the typical complex waveform (Figure 4A) also observed in the adult (Figure 1C). The FSL to monaural clicks decreased during development (Figure 4B). To obtain a measure of this decrease, we corrected the latencies for the cochlear delay and performed a linear regression on the juvenile data, yielding FSL = 5.59 ms – 0.064 ms/day (Figure 4C; *rho* = 0.39). By subtracting the delay between prespike and spike from the FSL, we measured the prespike latency (Figure 4D). This latency, after cochlear delay correction, also decreased during development (prespike latency = 4.59 ms – 0.051 ms/day, r = 0.33). As the FSL decreased more than the prespike latency, the prespike-AP delay decreased as well (Figure 4F, delay = 1.06 ms – 0.015 ms/day, r = 0.31), which was not due to a decrease in the EPSP-AP delay (Figure 4G, EPSP-AP delay = 0.42 ms – 0.003 ms/day, r = 0.19). Additionally, the AP-calcium channel activation and presynaptic release machinery may accelerate during this developmental period.

**Figure 4.**
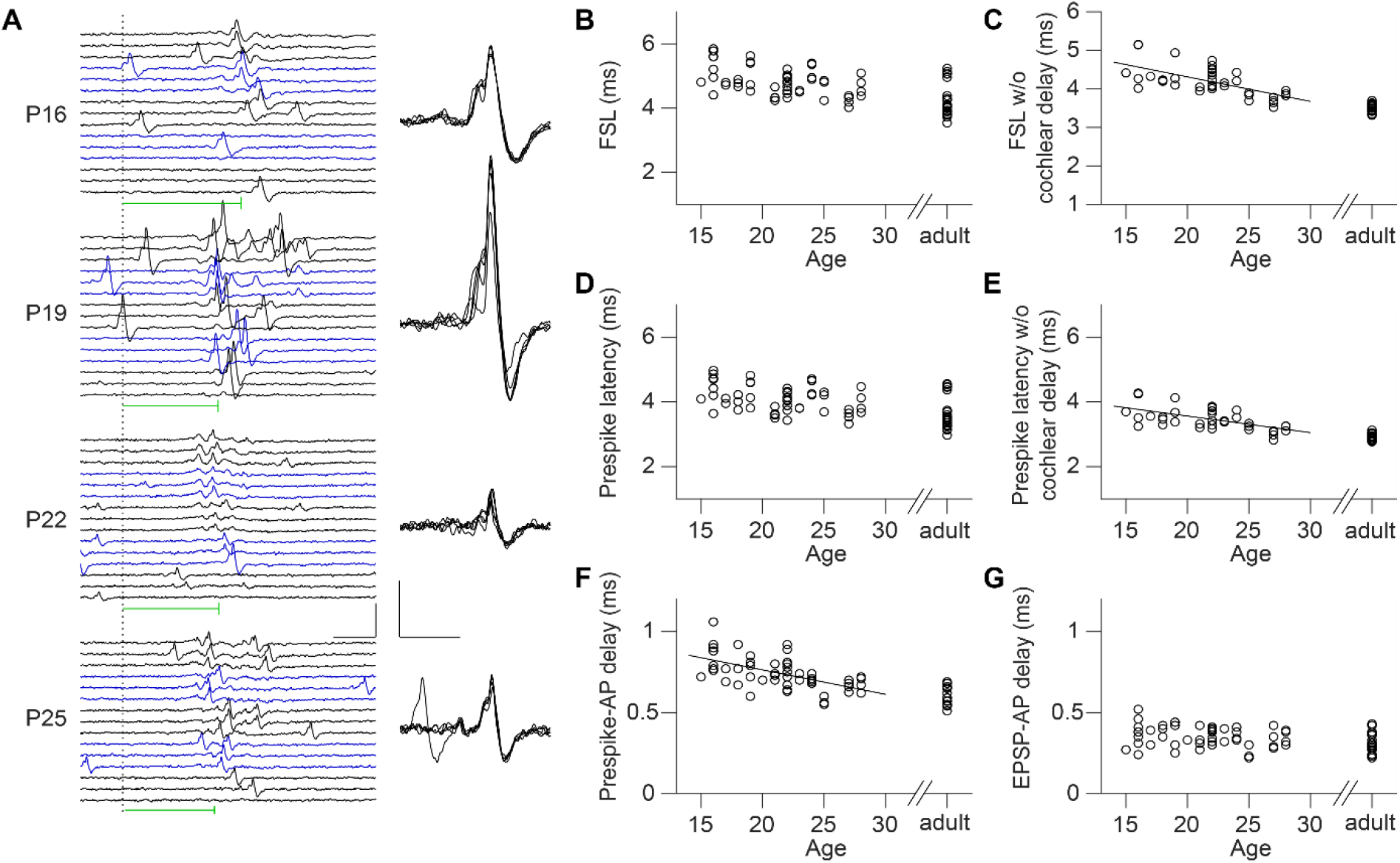
Bushy cell latencies decrease after hearing onset. (A) Left, Examples of responses to clicks of bushy cells following hearing onset. For each cell, responses elicited by clicks of increasing intensity (bottom-top) are shown, with three examples per intensity. Vertical dotted line indicates click onset. Horizontal green line indicates median latency for the cell. Scale bar: 3 mV, 5 ms. Right, five overlaid complex waveforms for each cell. Scale bar: 1 mV, 0.5 ms. (B) Bushy FSL against age. (C) FSL corrected for cochlear delay against age. (D-E) As in B-C for the prespike latency. (F) Prespike-AP delay against age. In B-G, each marker represents a bushy cell. In C, E and F, the line indicates the linear fit.

By extrapolation we can predict at what age the development of timing of the bushy cells is mature. All values (n = 17 adult bushy cells) in this section were corrected for the adult cochlear delay. The FSL was 3.53 ± 0.10 ms, which would be attained at 32 days after hearing onset. The prespike latency was 2.93 ± 0.10 ms, also attained at 32 days after hearing onset. Lastly, the prespike-AP delay was 0.60 ± 0.05 ms in the adult, estimated to be attained at 31 days after hearing onset.

Next, we studied developmental changes in the timing of ongoing activity using the Zwuis. Phase-locking of young BCs did not improve (Figure 5A-B), but a slight, non-significant increase in maximal firing rate was observed (Figure 5C). SdF of young low-frequency bushy cells was correlated with CF (r = 0.75, sdF = 0.43 kHz + 0.7 CF, n = 24 cells), excluding two cells with a CF of 5 kHz. While these two cells had an intensity-related change in delay of 0.55 and 0.60 ms, the delay changed only 0.05-0.32 ms for the other cells similar to adult bushy cells. As observed before (Englitz et al., 2009; Dehmel et al., 2010; Kuenzel et al., 2011; Nerlich et al., 2014; Keine and Rübsamen, 2015), young bushy cells were also resistant to synaptic depression (Figure 5D-G). Even at interspike intervals <5 ms and even for the youngest bushy cells we recorded from, the EPSP-AP delay was unaffected (Figure 5H). Young bushy cells thus encode the ongoing phase of their CF tone within complex sound accurately from hearing onset onward.

**Figure 5.**
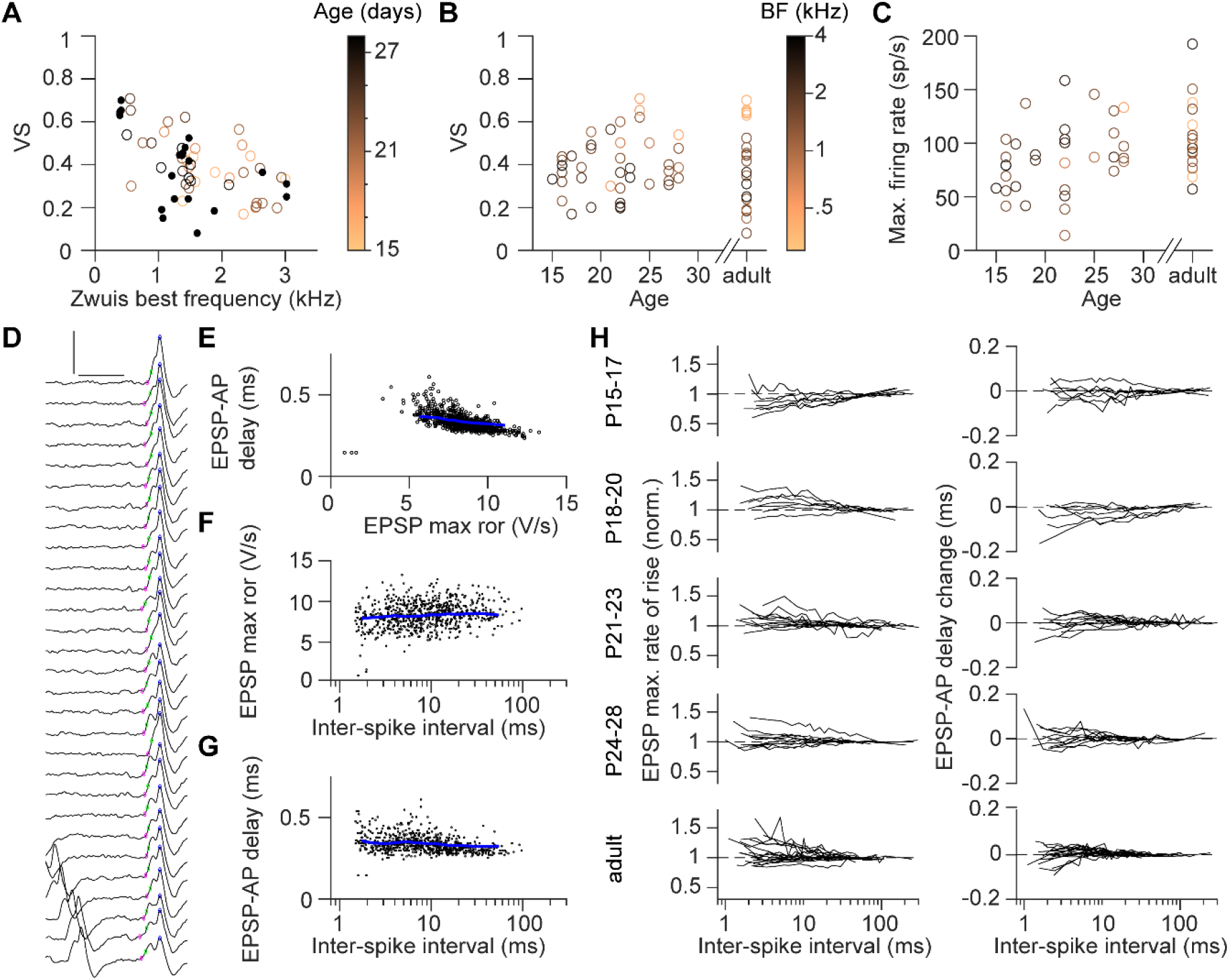
Timing of ongoing activity of developing bushy cells is already mature. (A) Vector strength (VS) against Zwuis best frequency. Age of the animal is color-coded, adult bushy cells are indicated by closed markers. (B) Vector strength against age. Zwuis best frequency is color-coded. (C) Maximal firing rate during Zwuis presentation against age. Zwuis best frequency is color-coded as in B. (D) Example traces aligned on spike peak and ordered from long-to-short interspike intervals of P23 bushy cell. Scale bar: 2 mV, 1 ms. Data in E-G are from the same cell. (E) EPSP-AP delay against EPSP max rate of rise. (F) EPSP max rate of rise against inter-spike interval. (G) EPSP-AP delay against inter-spike interval. For E-G, each black dot corresponds to a spike, blue line is the local average. (H) Normalized average EPSP max rate of rise (left) and the change in EPSP-AP delay (right) against inter-spike interval, separated by age. Each line corresponds to a bushy cell.

### Dissection of the axonal delay

With our analysis of bushy cell latency during development, we can pinpoint the contribution of the different neural delays to the decrease in latencies at the MSO. First, we subtracted the cochlear delay from the latencies of the developing MSO (Sierksma and Borst, 2024). After this correction, a developmental decrease in latencies was still there (Figure 6A, iMSO-FSL = 6.91 ms – 0.072 ms/day, rho = -0.68; cMSO-FSL = 7.63 ms – 0.096 ms/day, rho = -0.75). The adult latencies were 4.57 ± 0.04 ms and 4.63 ± 0.04 ms. This decrease in latency is partly explained by a decrease in EPSP-AP delay (Sierksma and Borst, 2024), which we fitted with a power function (*rho* = -0.6, p = 1 10^-5^, EPSP-AP latency = 0.461 ms + 161 age^-2.6^). What is not known is the synaptic delay from the arrival of the AP to the EPSP onset, but we will assume 0.4 ms, consistent with synaptic delays inferred from MSO’s field potentials (Clark and Dunlop, 1968) and with reported fast synaptic delays at other synapses (Borst et al., 1995; Sabatini and Regehr, 1996; Ritzau-Jost et al., 2014). In Figure 6D and Table 1 we summarized our findings on the development of latencies. Interestingly, the developmental decrease was not equal for the cMSO and iMSO. To isolate the developmental effects, we subtracted the adult latencies and plotted the change in latencies during development (Figure 6E). The developmental change is largely explained by a decrease in ear-BC prespike latency. For the cMSO-FSL we observed a decrease of 1.56 ms: 57% of this decrease is due to the ear-to-prespike latency (0.88 ms) and 24% due to a decrease in synaptic latencies (BC prespike-AP: 0.22 ms, MSO EPSP-AP latency: 0.16 ms). The remaining 19% or 0.30 ms was due to a decrease in BC-cMSO propagation latency. In contrast to the BC-cMSO, the BC-iMSO propagation latency did not decrease (Figure 6E C F). While the synaptic delay dominated the delay between BC and MSO, the developmental effect mostly arose from the propagation latency, which differentially decreased for iMSO and cMSO.

**Figure 6.**
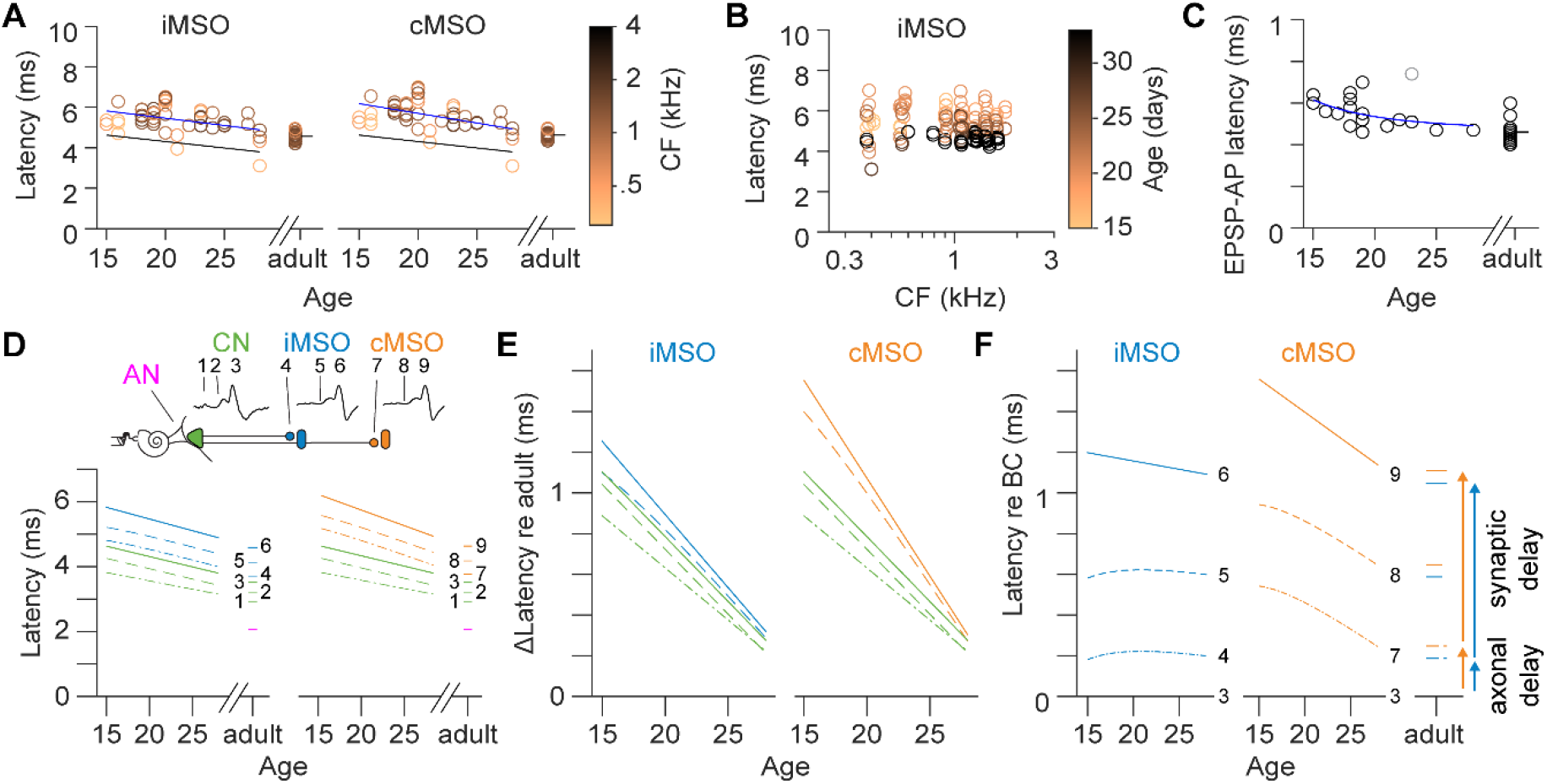
Dissection of the developmental decrease in latencies. (A) Latency of first spikes in ipsilateral and contralateral MSO (iMSO, cMSO) against age, corrected for cochlear delay. Each marker corresponds to an MSO neuron. Zwuis best frequency is color-coded. Blue line is linear regression fit. Black line is fitted FSL of bushy cells. (B) Latency of MSO neurons against best Zwuis frequency, corrected for cochlear delay. Age is color-coded. (C) EPSP-AP latency of MSO neurons against age. Data were fitted with a power function, excluding one data point (grey maker). (D) Summary of latencies across development. Schematic drawing indicates the legend of the panel. Solid lines denote FSL (3, c, S), dashed lines EPSP onset (2, 5, 8) and dash-dot lines presynaptic AP arrival (4, 7) or prespike (1). (E) Decrease in latency relative to adult latencies (Δ Latency) against age. Colors and line styles are as in D. (F) Latencies relative to BC-FSL against age. Colors, line styles and numbering are as in D.

**Table 1.** Latencies during development. Average latency and standard errors are listed. Values are corrected for the cochlear delay. For comparison, the ecological range of ITDs of adult Mongolian gerbils is −0.13 to +0.13 ms (Maki and Furukawa, 2005).

|  |  | **P15-21** |  |  | **P22-28** |  |  | **Adult** |  |  |
|---|---|---|---|---|---|---|---|---|---|---|
|  |  | mean | SE | n | mean | SE | n | mean | SE | n |
| **BC** | prespike (ms) | 3.61 | 0.36 | 14 | 3.33 | 0.32 | 21 | 2.93 | 0.10 | 17 |
|  | EPSP onset (ms) | 4.03 | 0.39 | 14 | 3.76 | 0.34 | 21 | 3.21 | 0.12 | 17 |
|  | FSL (ms) | 4.38 | 0.40 | 14 | 4.12 | 0.35 | 21 | 3.53 | 0.10 | 17 |
| **iMSO** | presyn (ms) | 4.43 | 0.20 | 5 | 4.35 | 0.25 | 3 | 3.65 | 0.20 | 6 |
|  | EPSP onset | 4.83 | 0.63 | 5 | 4.75 | 0.25 | 3 | 4.05 | 0.20 | 6 |
|  | FSL (ms) | 5.58 | 0.23 | 28 | 5.36 | 0.31 | 14 | 4.57 | 0.19 | 20 |
| **cMSO** | presyn (ms) | 4.61 | 0.23 | 5 | 4.53 | 0.27 | 3 | 3.74 | 0.17 | 6 |
|  | EPSP onset | 5.01 | 0.23 | 5 | 4.93 | 0.27 | 3 | 4.14 | 0.17 | 6 |
|  | FSL (ms) | 5.88 | 0.66 | 28 | 5.45 | 0.33 | 14 | 4.63 | 0.17 | 20 |

### Development of inter-nuclei distances and conduction velocity

Next, we studied the developmental changes in the auditory brainstem anatomy by reconstructing the AVCNs, MSOs and the MSO somatic layer at different developmental ages for 15 young and 7 adult gerbils (Figure 7A). The auditory brainstem development is illustrated in Figure 7B. Volumes of AVCN and MSO increased during this period (Figure 7C, AVCN: *rho* = 0.77, *p* = 2 10^-14^, Vol_AVCN_ = 335 nL + 315 nL (1+exp(22 days-age))^-1^, MSO: *rho* = 0.62, *p* = 2 10^-14^, Vol_MSO_ = 125 nL + 118 nL (1+exp((21 days-age)/0.6 days)^-1^). The approximate doubling of AVCN volume was in agreement with an earlier study (Gleich et al., 1998). Growth mainly occurred along the rostrocaudal axis (*rho* = 0.66 and 0.67 for AVCN and MSO, respectively), but not along the ventrodorsal (rho = 0.39 and -0.19 for AVCN and MSO, respectively) or mediolateral axes (rho = 0.10 and 0.30 for CN and MSO, respectively; Figure 7D). Consequently, the inter-nuclei distances remained relatively similar during development. The distance from AVCN to iMSO increased by ∼0.3 mm (*rho* = 0.78, young: d_AVCN-MSO_ = 1.94 mm + 0.016 mm/day age, adult: 2.58 ± 0.19 mm), while the inter-MSO distance is constant after hearing onset (rho = -0.16, young: 1.80 ± 0.09 mm, adult: 1.84 ± 0.12 mm, Figure 7E). The auditory brainstem thus grows rostrocaudally during the second week after hearing onset without affecting the distance between MSO nuclei.

**Figure 7.**
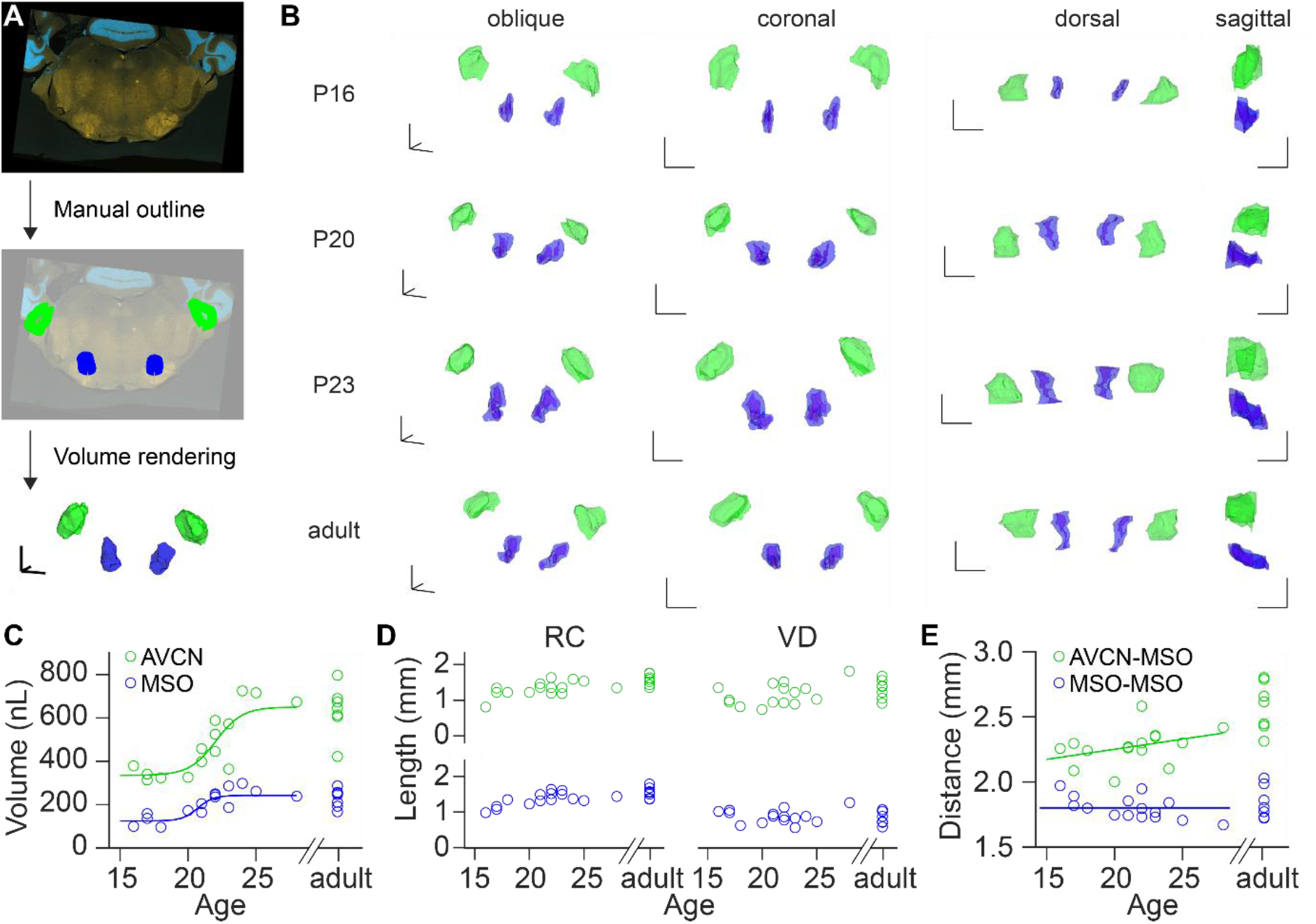
Auditory brainstem after hearing onset mainly grows in the rostrocaudal direction. (A) Brain sections were immunolabeled for VGluT1/2 and/or GlyT2 (top), AVCN (green) and MSO (blue) were manually outlined (middle) and rendered as volumes (bottom). (B) Four examples of 3D rendered nuclear volumes. From top to bottom, brainstems of P1c, P20, P23 and an adult gerbil. From left to right, an oblique view, a coronal view, a dorsal view and a sagittal view of the brainstem. Scale bars of 1 mm point toward dorsal, rostral, and medial. (C) Average volume of the AVCN (green) and MSO (blue) against age. (D) Bounding box length of AVCN (green) and MSO (blue) in the rostrocaudal (left) and ventrodorsal direction (right). (E) Average distance between AVCN and ipsilateral MSO (green), and between MSO and MSO (blue) against age.

The minor changes in the anatomical distances are contrary to the change in propagation latency, suggesting that the conduction velocity increases during this developmental period (Seidl and Rubel, 2016; Sinclair et al., 2017). To estimate average velocities, we combined the anatomical distances and *in vivo* latencies. In Figure 8A we plotted the distances against the average latency and the latencies against the average distance to visualize the spread in both dimensions. The angle of the fitted line corresponds to the average velocity of the bushy axon to the MSO. Clearly, the angle to the contralateral MSO is higher than to the ipsilateral MSO, indicating that the average propagation velocity to the contralateral MSO is higher. Velocities were increasing during development (Figure 8B-C).

**Figure 8.**
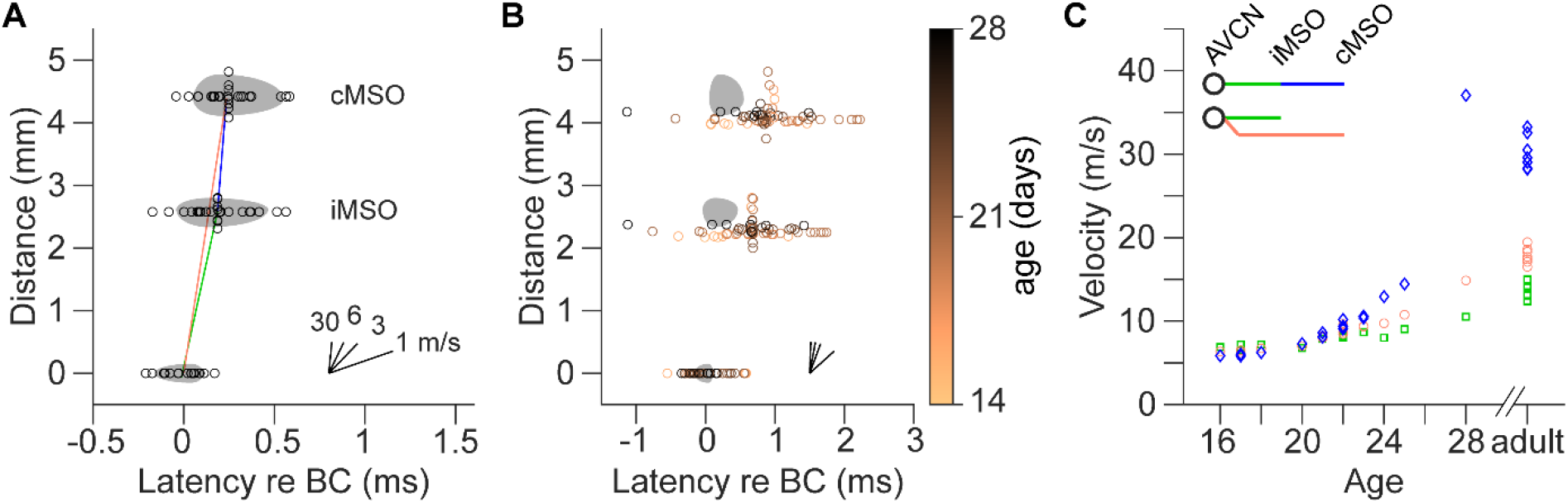
Differential increase in propagation velocity of the contralaterally projecting branch of the bushy axon. (A) Expected distance from AVCN to iMSO and cMSO against latencies. The distance between the AVCN and the cMSO is the sum of the AVCN-iMSO distance and the inter-MSO distance. Individual distances are plotted against average latency and average distances are plotted against individual latencies. The grey areas illustrate an ellipse that ranges from the 10^th^ to the S0^th^ percentile with its center of mass at the 50^th^ percentile. The angle corresponds to velocities as indicated by the black lines. The colored lines indicate the average velocities for the situations illustrated in panel C. (B) As in A, but the colored markers indicate the age of the animal. The grey areas pertain to the adult situation. (C) Velocity against age. The conduction velocity is calculated using the anatomical distance divided by the average latency. The velocity of the contralateral branch is calculated for two situations, as illustrated in the legend.

Bushy cells project to both MSOs (Osen, 1969; Smith et al., 1993; Beckius et al., 1999; Karino et al., 2011), but where the axon bifurcates into the ipsilateral and contralateral projecting axon is uncertain. We will consider two extremes here. At the one extreme, the contralateral branch already sprouts at the bushy cell and thus its latency/velocity is independent of the ipsilateral branch (CN-cMSO). At the other extreme, the axon branches at the ipsilateral MSO and thus the contralateral latency is equal to the ipsilateral latency plus the inter-MSO latency (MSO-MSO). The latter is closest to the anatomical situation, as bushy axons typically branch close to iMSO (Karino et al., 2011). The velocity for the contralateral branch can be estimated for either of these two extremes. After hearing onset, the mean velocities of CN-iMSO, CN-cMSO and MSO-MSO were 7.2 ± 0.5 m/s, 7.6 ± 1.2 m/s and 6.9 ± 1.1 m/s (P16-21). Both CN-cMSO and MSO-MSO velocities were not different from the ipsilateral velocity (paired T-test, *p* = 1.0 and 0.8). The adult CN-iMSO velocity, CN-cMSO velocity and MSO-MSO velocity increased to 14 ± 1 m/s, 18 ± 1 m/s and 30 ± 2 m/s, respectively (Figure 8C). For both extremes, the propagation velocity of the contralateral branch was higher than of the ipsilateral branch (paired T-test, *p* = 7 10^-7^ and 7 10^-7^). We conclude that the differential decrease in latency to the contralateral MSO is due to a higher velocity in the contralateral branch of the bushy axon and that this difference appears between P22-28.

### Variability in BC latencies underlies ITD tuning

Finally, we revisited a core assumption of the Jeffress’ model, namely that the axonal delays compensate for the ITD. Conceptually, in this model there are two bushy cells that both activate an MSO neuron, but are not activated by the same ear. The relative timing of bushy activation is solely determined by the ITD. Yet, we observed that there is intrinsic variation in FSLs across bushy cells. The variation in the averaged FSL ranged from 1.1 ms (P15-21), 0.8 ms (P22-28) to finally 0.26 ms in the adult (Table 1). The difference in the averaged FSL (ΔFSL) between bushy cells may partially compensate for ITDs. By calculating ΔFSL for all possible pairs, corrected for the cochlear delay (Figure 9), we found that the range of ΔFSL was larger than the ecological range. Moreover, bITDs are generally tuned to contralateral sound sources. To compare ΔFSL with bITDs, we added an age-specific contralateral delay—the difference in cMSO and iMSO latency—to ΔFSL (Figure 9). Although the distributions differed for group P15-21 (ks = 0.43, p < 0.001), they were not statistically different for group P22-28 and remarkably similar for the adult group. In principle, random pairing of bushy cells with the same CF can compensate adult ITDs.

**Figure 9.**
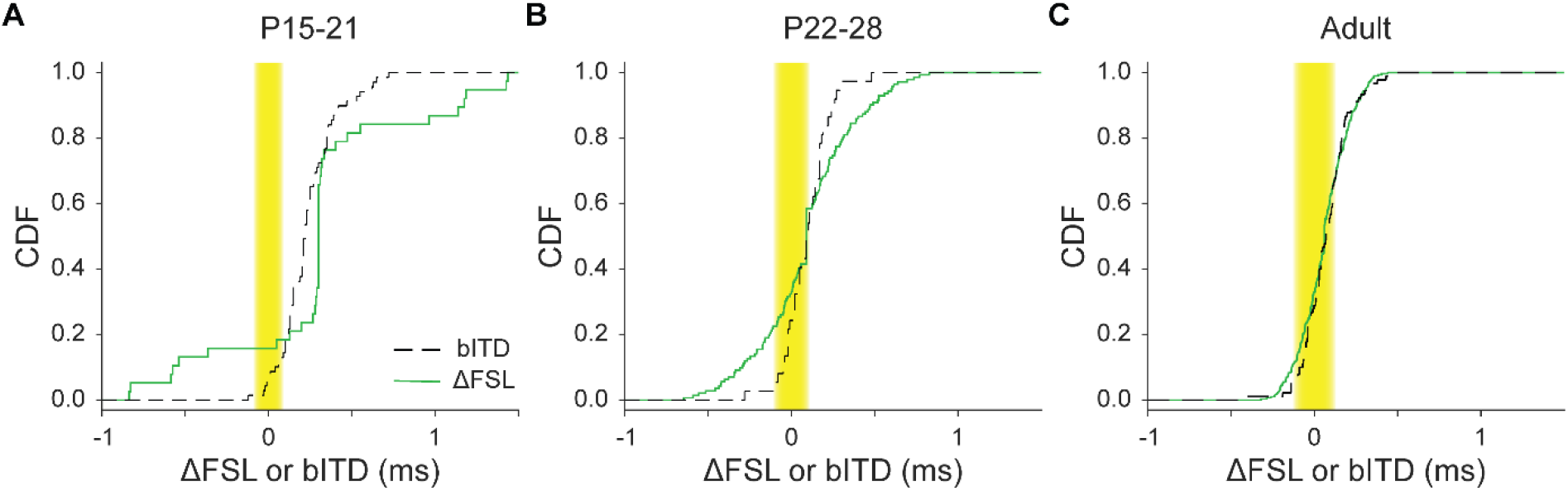
Random pairing of bushy latencies. (A) The cumulative distribution functions are shown for bushy FSL (ΔFSL) and bITDs for gerbils of P15 to P21. ΔFSL includes a contralateral delay (see Main text). Ecological range is indicated by the yellow shaded box. (B) As in A for gerbils of P22 to P28. (C) as in A for adult gerbils. Adult bITDs are taken from (Plauška et al., 2017; Sierksma and Borst, 2024) and young bITDs from (Sierksma and Borst, 2024).

## Discussion

We dissected the developmental changes in the auditory brainstem delays using *in vivo* recordings of bushy and MSO cells. Shortly after hearing onset BC firing was already precisely timed both for onset and ongoing complex sounds. Delays before the cochlear nucleus, between prespike and bushy cell spike, and to the MSO all decreased after hearing onset. We observed that the inter-MSO delay decreased without a similar decrease in the CN-iMSO axonal delay, suggesting a differential increase in propagation speed in the bushy axon branch crossing the midline. Finally, the differences in BC onset timing were sufficient to recreate the variation in bITDs of MSO neurons.

### Bushy and MSO neuron timing for tones vs complex sounds

Here, we measured the timing properties of bushy cells both in young and in adult gerbils, and we compared this to our earlier measurements on MSO neurons (add Sierksma and Borst, Plauska). For these comparisons we mostly used the responses to the wide-band stimulus Zwuis. In contrast to tones, wide-band stimuli (e.g. click, Zwuis, noise) elicit bushy output that is dominated by frequencies near CF, producing a stable delay (Louage et al., 2005; Wei et al., 2023)(this paper). In contrast, pure tones activate bushy cells with large and variable delays (Smith et al., 1991; Recio-Spinoso, 2012; Typlt et al., 2012; Wei et al., 2017). Below-CF, high intensity stimulation can elicit better phase-locking than at CF plus a phenomenon called peak-splitting, at which the neuron fires at more than one delay (Joris et al., 1994a; Wei et al., 2017; Wei et al., 2023), which may alter bITD (Batra et al., 1997; van der Heijden et al., 2013; Franken et al., 2015). Studies using tones reported bITDs significantly outside the ecological range (Spitzer and Semple, 1995; Pecka et al., 2008; van der Heijden et al., 2013; Franken et al., 2015), while studies using wideband stimuli typically find bITDs falling close to or within the ecological range (Plauška et al., 2016; Plauška et al., 2017; Mackenbach and Borst, 2023; Sierksma and Borst, 2024). The precise timing of the recruited bushy cells thus differs between complex sound and tones, which is important when dissecting the delays that determine the bITDs.

### Early maturation of bushy cells

We previously observed that the MSO’s ITD tuning at hearing onset is already remarkably mature (Sierksma and Borst, 2024). In agreement with this, in our present study we observed that many properties of the bushy cells were also already adult-like, including maximum firing rate and the presence of a complex waveform, sound level timing independence, phase locking, and resistance to synaptic depression.

### Dissection of latencies underlying ITD tuning

Directly after hearing onset the bITDs of the MSO neurons are mostly outside the ecological range (Sierksma and Borst, 2024). In the next two weeks neural latencies show a rapid decrease (Sierksma and Borst, 2024; Anbuhl et al., 2025). Combining our earlier MSO measurements here with bushy cell measurements we observed a preferential shortening of BC-cMSO axonal delays with little change in BC-iMSO axonal delays during development. Two factors contributed to this differential change. One is that the anatomical AVCN-MSO distance increased developmentally, whereas the distance between both MSOs remained the same. The second is that the conduction speed increased more for the contralateral projection than for the ipsilateral. There are several caveats about the latter conclusion. Firstly, because of our limited number of recordings from these BCs, we corrected their latencies by the estimated cochlear delay. Even though a similar cochlear delay was observed in the MSO latencies, we have not directly measured the latencies and the variation of BC with CF of 0.5-1 kHz, which are the primary input to the MSO (Grothe et al., 2010). Secondly, while we attribute the developmental decrease in contralateral delay to an increase in propagation velocity of the contralateral branch (Seidl and Rubel, 2016; Sinclair et al., 2017), without axonal reconstructions it cannot be excluded that the reported inter-nuclear distances differ from true axonal lengths. Bushy axons arborize extensively at the MSO (Beckius et al., 1999; Karino et al., 2011), and the additional length indicates that the inter-nuclear distance may underestimates the axonal length. On the other hand, axonal reconstructions indicate a smaller distance between AVCN-iMSO (1.94 vs. 1.40 mm) and a larger distance between MSOs (1.80 vs 2.1 mm,(Seidl and Rubel, 2016)). It remains uncertain how the reported BC-MSO delay arises from the axonal branches. Nevertheless, the combination of the stable BC-iMSO delay and decreasing BC-cMSO delay with the growth of the brainstem supports an increase in conduction velocity during this period (Seidl and Rubel, 2016; Sinclair et al., 2017).

Our reported conduction velocity of the contralateral branch (18-30 m/s) is higher than 3.6-11.0 m/s but still consistent with the highest reported latencies of 22-32.2 m/s (Clark and Dunlop, 1968). However, anatomical support for this higher contralateral velocity is limited as some found that the contralateral branch has a larger diameter (Smith et al., 1993; Seidl and Rubel, 2016) and longer inter-nodal distance (Seidl and Rubel, 2016) than the ipsilateral branch, while others did not find a difference in diameter (Beckius et al., 1999; Karino et al., 2011). Axon diameters are 1-3 μm (Brownell, 1975; Beckius et al., 1999; Karino et al., 2011; Ford et al., 2015; Sinclair et al., 2017), corresponding to 6-18 m/s (assuming 6 m/s per μm diameter in myelinated axons (Debanne et al., 2011)) consistent with the ipsilateral but not the contralateral velocity.

The decrease in latency may be ascribed to the delays in the terminal arborization instead of the main trunk. The profuse branching (Smith et al., 1993; Beckius et al., 1999; Karino et al., 2011) can be independently tuned at each MSO, and they create a range of differential delays because of the additional length and their lower conduction velocity. Shorter branches and/or faster conduction at the contralateral relative to ipsilateral branches can differentially reduce the latency. As another alternative explanation, MSO neurons may selectively strengthen inputs with matching latencies (Sierksma and Borst, 2024). This process favors the ‘fastest’ contralateral inputs, reducing the contralateral latency. As a 0.3 ms reduction is needed, it cannot be physiologically attained by selecting synapses closer to the soma, but this may contribute (Winters et al., 2017; Callan et al., 2021). Selecting axonal branches can create a 0.3 ms reduction in gerbils (Seidl and Rubel, 2016) or cats (Beckius et al., 1999; Karino et al., 2011). In either case, the differential delay then arises not in the main axonal trunk but at the final branches.

### Jeffress’ model revisited

The Jeffress’ model for ITD tuning implements MSO’s ITD sensitivity by combining a differential axonal delay with a coincidence detector, allowing them to be tuned to a specific sound direction. Generally speaking, our data are consistent with this model: neural delays seem to underlie the neuron’s bITD (Sierksma and Borst, 2024) and axonal delays can compensate for the ecological range of ITDs (Seidl and Rubel, 2016)(this paper). However, the intrinsic variability in BC timing was not considered before, whereas this creates a second possible source of internal delays. We compared the variance in the averaged BC delay with the variance in the bITD of the MSO neurons. Shortly after hearing onset, the variance in the BC delays greatly exceeded the variance in the bITDs, whereas in the adult, both distributions were closely matched. The young MSO neuron may be contacted by 7-70 inputs (Callan et al., 2021) which is further refined to 4-8 inputs (Couchman et al., 2010). We previously used a simulation to show that this larger pool of inputs at hearing onset reduces the variation in bITDs and centers the bITDs at the averaged contralateral delay (Sierksma and Borst, 2024). The excess variance of BC delays in young gerbils suggests the presence of a pruning mechanism that selects for matching BC delays, which would improve the sharpness of ITD tuning.

Our data thus shed light on the remarkably precise mechanisms that allow gerbils to localize sounds along the azimuth. Each MSO is biased to contralateral sound sources. Shortly after hearing onset the bITDs lie outside of the ecological range, and firing will be maximal for contralateral sound, allowing the MSOs to already encode direction (“slope coding”; (McAlpine et al., 2001)). The developmental changes bring the bITDs toward the ecological range, but a contralateral bias remains close to the ecological limit of the gerbil, which most likely derives from the inter-MSO delay imposed by the contralateral branch (0.1 ms = 1.8 mm / 18 m/s; (Maki and Furukawa, 2005; Sierksma and Borst, 2024). Despite its small head size, it is thus able to read out the minute differences in arrival time of sounds at both ears efficiently.

## Acknowledgments

This work was financially supported by the Nederlandse Organisatie voor Wetenschappelijk Onderzoek (NWO; ALW-open, ‘Great Timing’, #824.15.008 to JGGB; ENW-XS, ‘All ears’, OCENW.XS22.151 to MCS and ENW-M, ‘Learning to localize sounds’, OCENW.M.21.218 to JGGB). The funding agency had no role in study design, data collection and analysis, decision to publish, or preparation of the manuscript. We thank Dr. Marcel van der Heijden for providing the software for sound stimulus generation and analyses, and Dr Aaron Wong for helpful discussions.

## Author Contributions

MCS designed research, performed research, analyzed data, wrote the first draft of the paper, wrote the paper. JGGB designed research, wrote the paper, edited the paper.

